# Natural selection shapes codon usage in the human genome

**DOI:** 10.1101/688051

**Authors:** Ryan S Dhindsa, Brett R Copeland, Anthony M Mustoe, David B Goldstein

## Abstract

Synonymous codon usage has been identified as an important determinant of translational efficiency and mRNA stability in model organisms and human cell lines. However, to date, population genetics studies have failed to observe evolutionary constraint on human codon usage, and synonymous variation has been largely overlooked as a component of human genetic diversity. Using genetic sequencing data from nearly 200,000 individuals, we uncover clear evidence that natural selection optimizes codon content in the human genome. We derive intolerance metrics to quantify gene-level constraint on synonymous variation and demonstrate that dosage-sensitive, DNA damage response, and cell cycle regulated genes are more intolerant to synonymous variation than other genes in the genome. Notably, we illustrate that reductions in codon optimality can attenuate the function of BRCA1. Our results reveal that synonymous mutations likely play an important and underappreciated role in human variation.

## Introduction

A long-standing assumption in human genetics has been that synonymous mutations are unimportant because they do not alter the resulting protein sequence. In fact, the synonymous mutation rate is typically designated as the background neutral mutation rate against which other forms of mutational selection is measured (Kimura, 1977), and synonymous mutations are almost always ignored in genetic disease association studies. However, recent evidence indicates that synonymous variation is not always neutral and may often have functional consequences (Hanson and Coller, 2018; Hunt et al., 2014).

Well-established mechanisms through which synonymous mutations can impact gene expression include disruption of splicing enhancer sites (Parmley et al., 2006; Supek et al., 2014), mRNA secondary structure (Shen et al., 1999), and binding sites for regulatory RNA-binding proteins and microRNAs (Brest et al., 2011; Capon et al., 2004). Significant progress has been made in identifying these “functional” synonymous mutations due to overlap of known regulatory sites and proximity to exon boundaries. Although much less understood, emerging evidence suggests that synonymous mutations in the gene body can also impact gene expression. Specifically, biochemical studies indicate that “optimal” codons matching more abundant tRNAs can support rapid translation, whereas synonymous but “non-optimal” codons can slow translation (Bazzini et al., 2016; Forrest et al., 2018; Presnyak et al., 2015; Wu et al., 2019). Importantly, synonymous codon usage also seems to affect human mRNA stability via coupling between mRNA degradation and translation (Forrest et al., 2018; Wu et al., 2019). Indeed, it has long been recognized that the human genome exhibits clear codon usage biases, with certain codons used more frequently than others (Chamary et al., 2006; Eyre-Walker, 1991). Nevertheless, the significance of human codon usage bias as it relates to human physiology and fitness remains controversial.

Efforts to detect selective pressures on codon usage in the human genome have been hampered by three main challenges. First, synonymous mutations are expected to be weakly deleterious, as they are more likely to affect protein abundance than function (Chamary et al., 2006; Keightley and Eyre-Walker, 2000). Because the small effective population size of humans limits the efficacy of natural selection in purging weakly deleterious mutations, any evolutionary constraint on synonymous mutations will only be evident with a large sample size of sequenced individuals. Second, codon usage is posited to be functionally linked to tRNA expression (Gingold et al., 2014; Wu et al., 2019). Because tRNA expression varies widely by tissue (Dittmar et al., 2006), synonymous sites are likely subject to competing pressures that convolute the evolutionary signature. This variation in tRNA expression also makes it difficult to correlate codon usage with tRNA availability. Third, the nucleotide content at synonymous sites strongly correlates with local GC content in nearby non-coding regions. This phenomenon suggests that codon bias is influenced by evolutionarily neutral processes, such as local variation in mutation rate (Chamary et al., 2006; Duret and Mouchiroud, 1999; Eyre-Walker, 1991; Hershberg and Petrov, 2008; Pouyet et al., 2017). These challenges have led to conflicting data, with much of the field believing codon bias is a passenger effect rather than a result of selective forces (Chamary et al., 2006). Altogether, these challenges necessitate a robust statistical method that can detect selective constraint on variants of modest effect across a population while controlling for confounding mutational biases.

In this study, we leveraged the unprecedented amount of sequencing data available in two population reference cohorts — TOPMed (62,784 genomes) (Taliun et al., 2019) and gnomAD (123,136 exomes) (Karczewski et al., 2019) — to demonstrate that natural selection optimizes codon content in protein-coding regions in the human genome. We devised two scores to rank genes by their intolerance to synonymous mutations. The first metric, synRVIS, measures human-specific constraint against changes in codon optimality. The second metric, synGERP, reflects phylogenetic conservation at fourfold degenerate sites across the mammalian lineage. These scores, in turn, allow us to identify genes and pathways in which synonymous variants are most likely to affect human fitness. Overall, our work provides the first direct genome-wide evidence that codon usage in humans is under natural selection and has strong implications for the interpretation of synonymous variants in the human genome.

## Results

### Site frequency spectra reveal genome-wide signatures of purifying selection on human codon usage

The availability of aggregated human whole-genome allele frequency data from roughly 60,000 individuals contained in the TOPMed database (Taliun et al., 2019) provides an unprecedented resource for investigating selective constraint on weakly deleterious variants such as synonymous mutations. Focusing on synonymous sites where any of the four nucleotides in the third position of the codon encode the same amino acid (i.e. four-fold degenerate) synonymous sites, we sought to use this resource to identify potential evidence of natural selection on codon usage. The standard approach for measuring purifying selection is to examine the allele frequency spectrum. Allele frequency is a powerful proxy of a variant’s phenotypic impact, as purifying selection tends to eliminate deleterious variants before they reach a high frequency in the population (Lappalainen et al., 2019), and hence the spectrum should skew relative to the neutral mutation rate. Typically, the neutral rate is defined as the synonymous mutation rate. To enable robust comparisons, we generated a neutral reference set of variants by matching each observed synonymous variant to a nearby (<10 kilobases) randomly sampled intronic variant (**Fig S1A**). This procedure matches the GC content of the neutral reference to the synonymous test set, mitigating regional- and transcription-associated biases in mutation rates, and normalizing the total number of variants included in each set (Lawrie et al., 2013; Machado et al., 2017).

Following the classical approach, we first compared the site frequency spectrum (SFS; i.e. the distribution of allele frequencies) of synonymous variants and matched intronic variants (n=2,896,436) without accounting for changes in codon usage. Consistent with prior studies, the two distributions appeared nearly identical with the synonymous SFS exhibiting a very slight skew toward rarer allele frequencies (t-test *p* = .02) (**Fig S1B**). Thus, in aggregate, synonymous variants do not appear to be under significantly more constraint than putatively neutral intronic variants.

While aggregate analysis suggests that synonymous sites are not constrained, this analysis treats all synonymous variants as equivalent, ignoring that different variants may experience distinct selective pressures. Specifically, under the codon optimality hypothesis, a synonymous variant that increases codon optimality should be favored, whereas mutations away from optimal codons should be deleterious. While conceptually straightforward, the codon optimality hypothesis has remained challenging to test due to the challenge of classifying codon optimality. Unlike unicellular organisms, codon optimality cannot be matched to tRNA gene copy number because tRNA expression varies widely by tissue (Dittmar et al., 2006; Pan, 2018). Other common metrics, such as relative synonymous codon usage (RSCU), quantify codon frequencies (Sharp and Li, 1986), which can be confounded by many variables and do not directly reflect a codon’s effect on translational efficiency. We thus employed recently published codon stability coefficients (CSC), defined as the Pearson correlation coefficient between codon frequency in a transcript and the half-life of the transcript in human embryonic kidney 293T (HEK293T) cells. Although the molecular mechanism linking codon usage and mRNA stability in human cells remains an outstanding question, these scores moderately correlate with tRNA levels (Wu et al., 2019). Therefore, one proposed model is that human codon optimality partially reflects translation elongation speed (**Fig 1A**), which has been previously shown to affect mRNA stability in yeast (Radhakrishnan et al., 2016).

**Figure 1.**
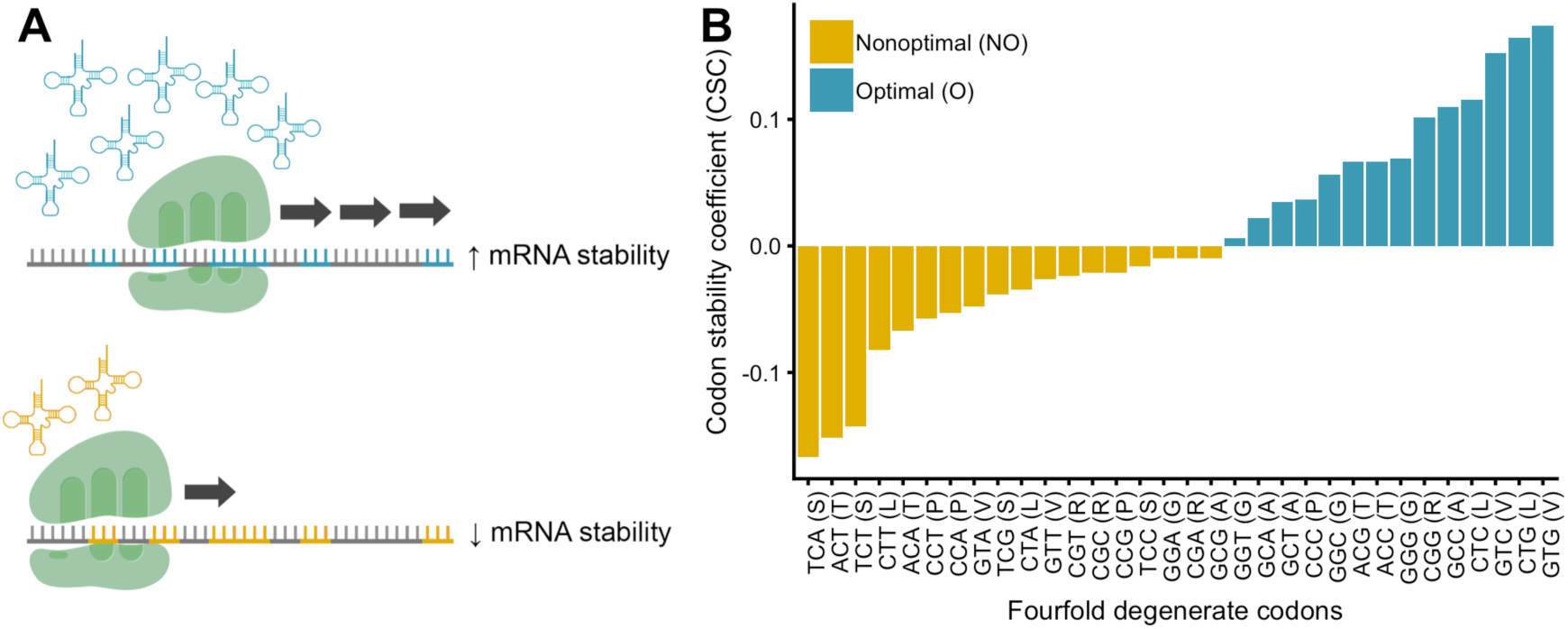
tRNA availability and codon usage affect mRNA decay. **(A)** Codon-optimized transcripts (top) exhibit increased mRNA stability compared to less optimized transcripts (bottom). Codon optimality modestly correlates with tRNA availability, suggesting optimal codons are decoded by more highly expressed tRNAs and may therefore increase translational efficiency. **(B)** Distribution of codon stability coefficient scores for fourfold degenerate codons previously derived by Wu et al., 2019 from HEK293T cells and depiction of our classification of optimal and non-optimal codons.

We classified the synonymous variants that resulted in changes from a codon with a positive CSC to a negative CSC as “optimal-to-nonoptimal” (O → NO), the opposite as “nonoptimal-to-optimal” (NO → O), and all others as “neutral.” Strikingly, O → NO synonymous variants segregated at significantly lower frequencies than matched intronic variants (*p* = 3.3 × 10^−33^), neutral synonymous variants (*p* = 1.14 × 10^− 35^), and NO → O synonymous variants (*p* = 4.2 × 10^−88^) (**Fig 2A**), suggesting constraint against variants that reduce codon optimality. Furthermore, NO → O synonymous variants segregated at significantly higher allele frequencies than their matched intronic variants (*p* = 1.9 × 10^−14^) and neutral synonymous variants (*p* = 3.0 × 10^−32^) (**Fig 2A**), implicating a role of positive selection in optimizing codon content. Similar results were observed when controlling for trinucleotide context (**Fig. S2A, B**), further supporting that the NO → O and O → NO allele frequency differences cannot be explained by local mutation rate differences.

**Figure 2.**
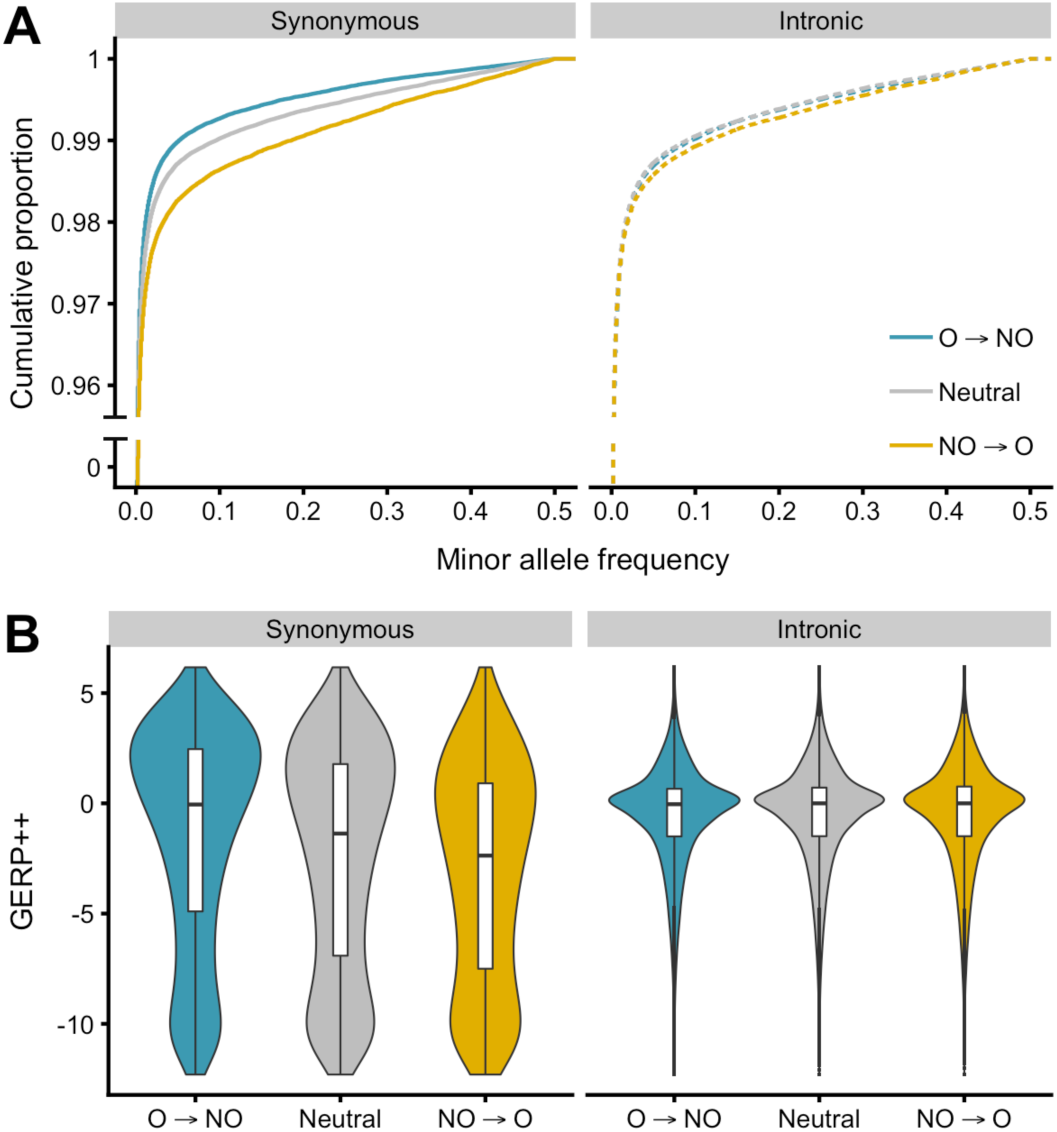
SFS and GERP++ distributions reflect selection on codon usage. **(A)** Site frequency spectra of synonymous variants that result in optimal-to-nonoptimal (O → NO), neutral, and nonoptimal-to-optimal codon (NO → O) changes (left) and matched intronic variants (right). **(B)** Distribution of GERP++ scores for the reference alleles of the variants included in (A).

The CSC scores were derived from mRNA stability measurements in a single cell line and therefore may not represent tissue-specific codon optimality patterns. We therefore repeated our SFS analysis using RSCU to define codon optimality instead of CSC. Notably, RSCU and CSC are significantly correlated (Pearson’s *r* = 0.41, *p* < 10^− 300^), indicating that optimal codons appear more frequently in the human genome. Furthermore, we observe similar evidence of purifying selection against O → NO variants and positive selection on NO → O variants when using RSCU to partition the SFS (**Fig S2D**).

Combined, our results implicate a role of negative selection in purging synonymous variants that reduce codon optimality and a role of positive selection in favoring variants that increase codon optimality. Despite the challenges associated with assessing codon usage in human cells, these findings strongly suggest that codon usage contributes to human genetic diversity and shapes human evolution.

### Optimal synonymous sites are evolutionarily conserved across the mammalian lineage

The shifts in the site frequency spectrum provide clear evidence of human-specific selective pressures on codon usage. Given that the link between codon optimality and translation should be general, we hypothesized that we should also observe signatures of conservation at fourfold degenerate synonymous sites across related phylogenetic species. To test this prediction, we assessed conservation using GERP++, a method that assigns each genomic position a score denoting its estimated evolutionary constraint across the mammalian lineage (Davydov et al., 2010).

Synonymous sites that are strongly conserved in the human genome have higher GERP++ scores than less conserved sites. Consistent with the hypothesis that optimal codons experience stronger evolutionary conservation, the GERP++ scores of the reference sites of O → NO synonymous variants were significantly higher than at matched intronic sites (Mann Whitney U *p* = 1.2 × 10^−31^), neutral synonymous sites (*p* < 10^−300^), and NO → O synonymous sites (*p* < 10^−300^) (**Fig 2B**). Moreover, the GERP++ scores of the reference sites of NO → O variants were significantly lower than matched intronic (*p* < 10^−300^) and neutral synonymous sites (*p* < 10^−300^), suggesting weaker phylogenetic constraint at nonoptimal sites. The GERP++ distributions of trinucleotide-matched and RSCU-annotated codon changes corroborated this observation (**Fig S2C, S2E**). Whereas the prior analysis only considered sites that were variant in the TopMED cohort, we next considered every fourfold degenerate site in the coding genome, and found that GERP++ significantly correlated with both CSC (Pearson’s *r* = .26, *p* < 10^−300^) and RSCU (*r* = .25, *p* < 10^−300^). Together, these results demonstrate long-term evolutionary pressures on codon usage, challenging prior assumptions that the small effective population sizes of mammals preclude selection from acting on synonymous sites.

### Human genes display intergenic variation in intolerance to synonymous variation

Our observations illustrate genome-wide signatures of constraint on codon optimality. However, we suspected that synonymous variation might be under stronger selective constraint in some genes than others. Therefore, we sought to quantify the strength of selection on synonymous sites per gene. We previously introduced the residual variation intolerance score (RVIS)—a scoring system that quantifies per-gene intolerance to functional (i.e. missense and loss-of-function) mutations using standing human variation (Petrovski et al., 2013). Here, we extended this framework, in an approach we term synonymous RVIS (synRVIS), to quantify genic constraint against synonymous variants that reduce codon optimality.

synRVIS only considers variation in the protein-coding genome. Therefore, to increase our sample size for constructing synRVIS, we used sequence data from the 123,136 exomes contained in gnomAD (Karczewski et al., 2019) rather than the roughly 60,000 genomes contained in TopMED. Specifically, we regress the number of common (MAF > 0.5%) O → NO synonymous variants on the total number of observed synonymous variants for each gene (**Fig 3A**). The resulting regression line predicts the expected number of common O → NO variants accounting for genic mutation rates, sequence context, and gene size. The deviation of each gene from this expectation (more or less variation than expected) is calculated as the studentized residual, with a synRVIS score below 0 indicating higher intolerance to O → NO synonymous variation.

**Figure 3.**
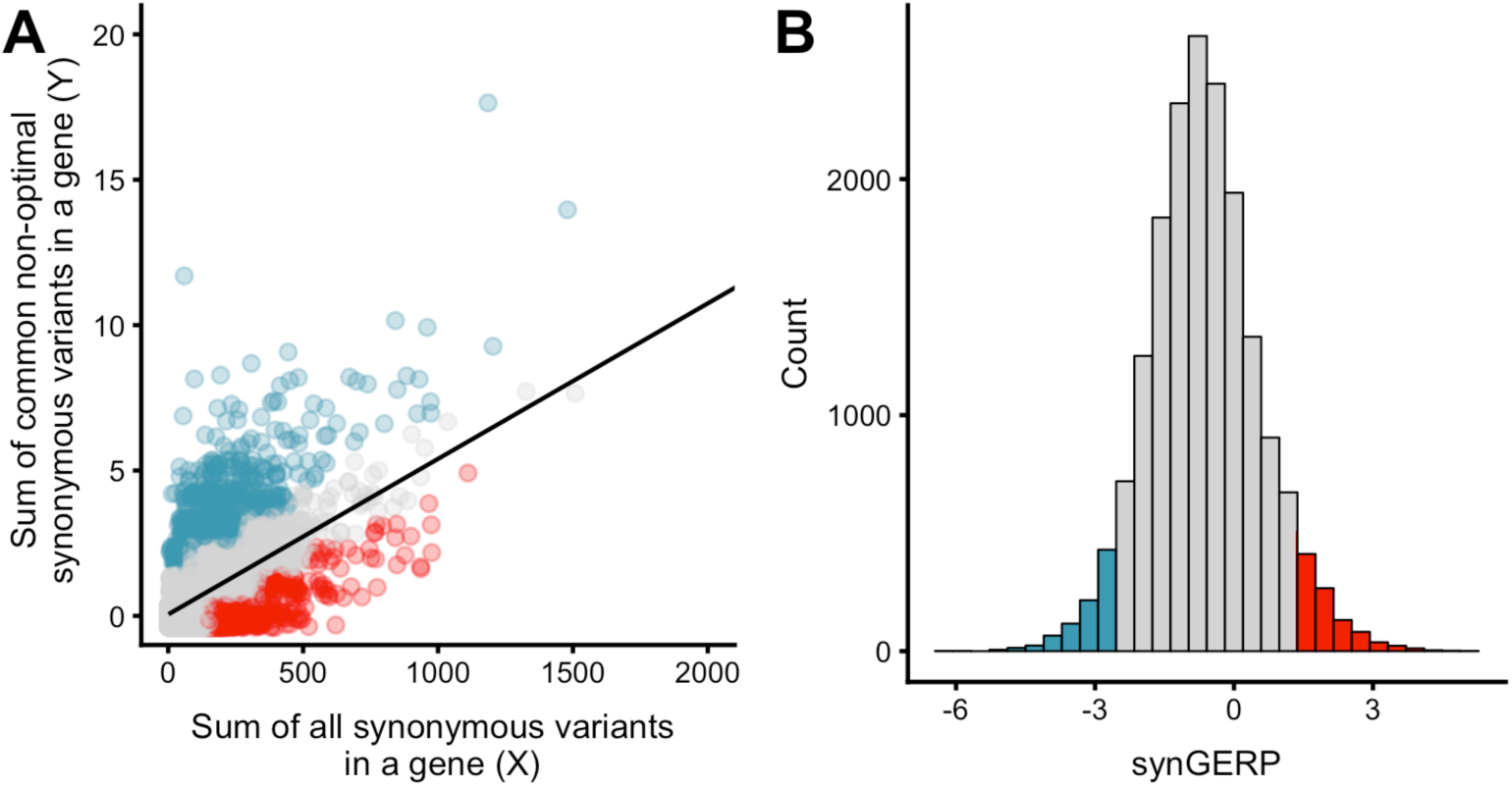
synRVIS derivation and distribution of synGERP scores. **(A)** synRVIS regression plot in which each point represents a gene. Red points represent the bottom fifth percentile (most intolerant) and blue points represent the upper fifth percentile (most tolerant). Two outlier genes with greater than 2,000 synonymous variants are excluded. **(B)** The distribution of synGERP scores. As in (A), color coding corresponds to the fifth percentile extremes.

synRVIS provides a direct, gene-specific measure of selection on codon optimality the human lineage. However, the dynamic range of the synRVIS metric is limited by the comparably small number of mutations at synonymous sites in gnomAD (median of 66 per gene). We therefore created a complementary score, which we termed synGERP, to quantify phylogenetic conservation at synonymous sites across the mammalian lineage. In order to create a per-gene metric, we took the mean GERP++ score at all fourfold degenerate synonymous sites in a given gene’s canonical transcript, excluding all codons adjacent to exon-intron boundaries. A higher synGERP score signifies overall stronger evolutionary conservation at fourfold degenerate sites for that gene (**Fig 3B**). To facilitate interpretation of these scores, we calculated genome-wide percentile scores for synRVIS and synGERP, in which a lower percentile indicates higher intolerance (synRVIS) or higher phylogenetic conservation (synGERP) (all scores are available in **Table S1**).

Interestingly, synRVIS and synGERP were only weakly correlated (*r*^2^ = .013, *p* = 2.3 × 10^−51^). We have similarly observed low correlation between human-specific intolerance scores and GERP-derived scores in prior evaluations of non-coding regulatory regions (Petrovski et al., 2015). One possible explanation for this low correlation is that a fraction of codon usage may be under human-specific selection, for example mirroring human-specific tRNA expression patterns, which would only be captured by synRVIS. Additionally, whereas synRVIS isolates codon optimality effects, synGERP measures the combined constraint on synonymous sites from sources such as splicing enhancers and RNA-binding protein binding sites. Together, these two scores provide a framework for identifying genes that are most intolerant to synonymous variation.

Gene ontology (GO) enrichment tests revealed that the most synRVIS intolerant genes (< 25^th^ percentile) were enriched for ontologies related to the cell cycle and transcription, including “cellular response to DNA damage,” “microtubule-based processes,” and “positive regulation of transcription by RNA polymerase II.” Furthermore, synGERP intolerant genes were enriched for ontologies such as “regulation of proteolysis involved in cellular protein catabolic process,” “regulation of mRNA stability,” and “negative regulation of translation” (**Table S3**). These results mirror observations in model organisms that the most codon optimized genes tend to be related to stress responses, translation, and post-transcriptional gene regulation (Burow et al., 2018; Carneiro et al., 2019) and therefore underscore the evolutionary significance of codon optimality.

### Genes intolerant to synonymous variation are enriched for dosage sensitive genes

Given the impact of codon usage on mRNA stability and protein expression, we hypothesized that well-known dosage-sensitive genes would be more intolerant to synonymous variation than other genes in the genome. To test this hypothesis, we constructed a logistic regression model to determine whether synRVIS and synGERP could predict the 360 genes denoted as dosage sensitive in ClinGen’s Genome Dosage Map (Rehm et al., 2015). We found that both synRVIS and synGERP significantly predicted this gene set: *p* = 8.2 × 10^−9^ (AUC = 0.60) and *p* = 2.2 × 10^−34^ (AUC = 0.68), respectively (**Fig 4A**). A joint model containing both scores achieved an AUC of 0.69, in which both synRVIS and synGERP remain predictive (*p* = 7.6 × 10^−7^ and *p* = 1.4 × 10^−31^), indicating that both scores provided significant independent information in predicting dosage sensitive genes.

**Figure 4.**
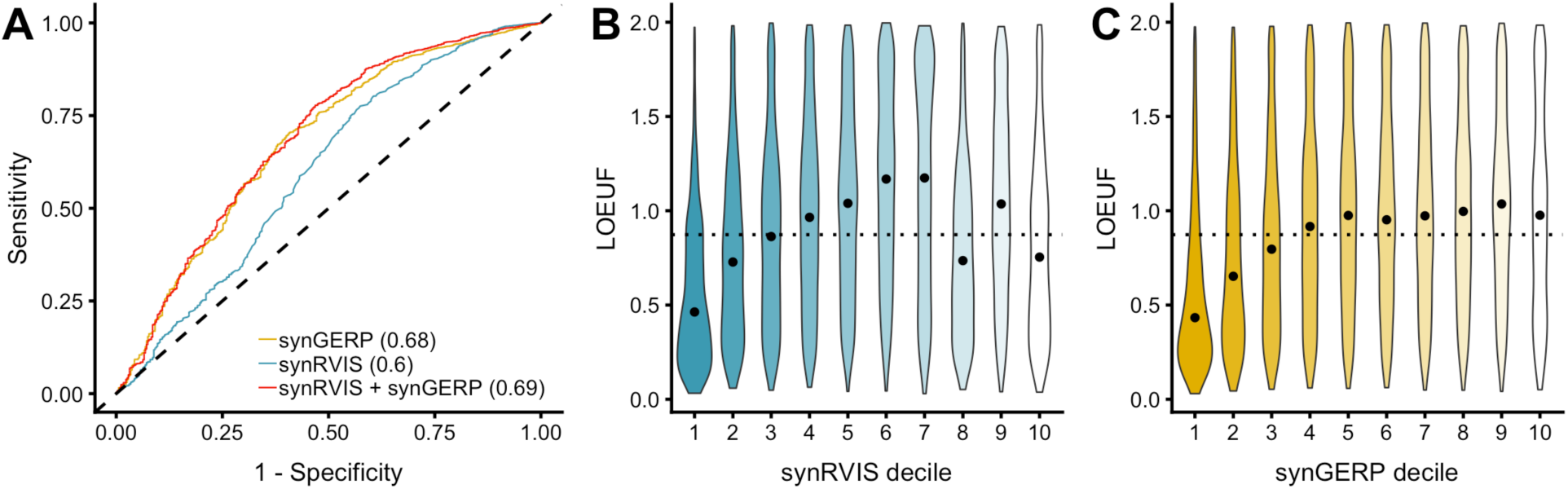
Dosage-sensitive genes are intolerant to synonymous variation. **(A)**. ROC curve demonstrating the capacity for synRVIS, synGERP, and a joint model to predict ClinGen dosage sensitive genes. AUCs for the respective models indicated in parentheses. **(B, C)** The distribution of LOEUF scores for each synRVIS and synGERP decile. The black dot indicates the median LOEUF score per synRVIS decile and the dotted horizontal line indicates the median LOEUF score across all genes.

The ClinGen dosage sensitive genes included in the prior analysis only include genes implicated in Mendelian disease. Another way to identify dosage sensitive genes is to identify genes depleted of loss-of-function variants in the human population. To verify that dosage sensitive genes are intolerant to synonymous variation, we compared synRVIS and synGERP to LOEUF (loss-of-function observed/expected upper bound fraction), a metric that represents the ratio of observed/expected loss-of-function variants within the gnomAD database (Karczewski et al., 2019). A lower LOEUF indicates higher constraint against loss-of-function variation. To compare synonymous constraint to LOEUF, we plotted the median LOEUF score per synRVIS and synGERP decile (**Fig 4B, C**). We observe that genes more intolerant to synonymous variation tend to be depleted of loss-of-function variation. Furthermore, synRVIS and synGERP both correlated with LOEUF (*r* = .15, *p* = 3.2 × 10^−89^ and *r* = .24, *p* = 3.3 × 10^−231^, respectively). However, we were surprised to find that some highly synRVIS tolerant genes were also enriched for low LOEUF scores (**Fig 4B**). This discordance implies that certain loss-of-function intolerant genes are tolerant to changes in codon usage.

A gene ontology (GO) enrichment analysis revealed that synRVIS-tolerant (>75^th^ percentile) but LOEUF-intolerant (<25^th^ percentile) genes were significantly enriched for certain neurodevelopmental pathways, such as “regulation of dendrite morphogenesis,” “positive regulation of axonogenesis,” and “synaptic vesicle endocytosis” (**Fig S4**). Notably, neurons are subject to different translational regulation programs than other cell types due to mTOR signaling (Blair et al., 2017) and their unique cellular demands, such as local translation at synapses (Holt and Schuman, 2013). Furthermore, recent evidence suggests that codon optimality may in fact be attenuated in the developing nervous system (Burow et al., 2018).

In sum, both synRVIS and synGERP can broadly predict dosage sensitive genes. These results emphasize the importance of codon usage in regulating gene expression and demonstrate that natural selection more strongly optimizes codon content in genes where differences in protein levels strongly impact human physiology.

### DNA damage genes and periodically expressed cell cycle genes are intolerant to changes in codon usage

If codon optimality is important in regulating gene expression, it is likely to not only be under particularly strong constraint in haploinsufficient genes, but also in genes that are expressed under limiting tRNA conditions. The cytoplasmic tRNA pool changes dynamically in terms of its overall abundance as well as its composition in response to cellular demands (Chan et al., 2012; Saikia et al., 2016; Torrent et al., 2018). We expected that genes that need to be highly expressed when tRNA levels are low should be the most intolerant to reductions in codon optimality.

Among the classes of stress response genes, we expected DNA damage repair genes to be under particularly strong constraint. In yeast, stress due to DNA damaging compounds results in reduced tRNA export from the nucleus as well as tRNA modifications that enhance translation of key DNA repair proteins (Begley et al., 2007; Ghavidel et al., 2007). In mice, knocking out the Elongator complex, which is required for translating codon-biased genes, leads to dysregulation of codon-biased DNA damage genes (Goffena et al., 2018). Motivated by these findings, we tested whether a previously published list of 178 DNA damage response genes were intolerant to synonymous variation (Wood et al., 2005). In a logistic regression model, synRVIS, but not synGERP, was able to predict genes involved in DNA damage response (AUC = 0.61, *p* = 6.02 × 10^− 05^; AUC = 0.52, *p* = 0.6, respectively) (**Fig 5A**). This result implies that codon usage in DNA damage repair genes is under human-specific constraint and thus likely plays an important role in regulating this pathway. Although our synGERP analysis suggests that codon bias is not conserved across eukaryotes, we suspect this discordance between synRIVS and synGERP is due to species-specific variation in the stress-induced tRNA pools.

**Figure 5.**
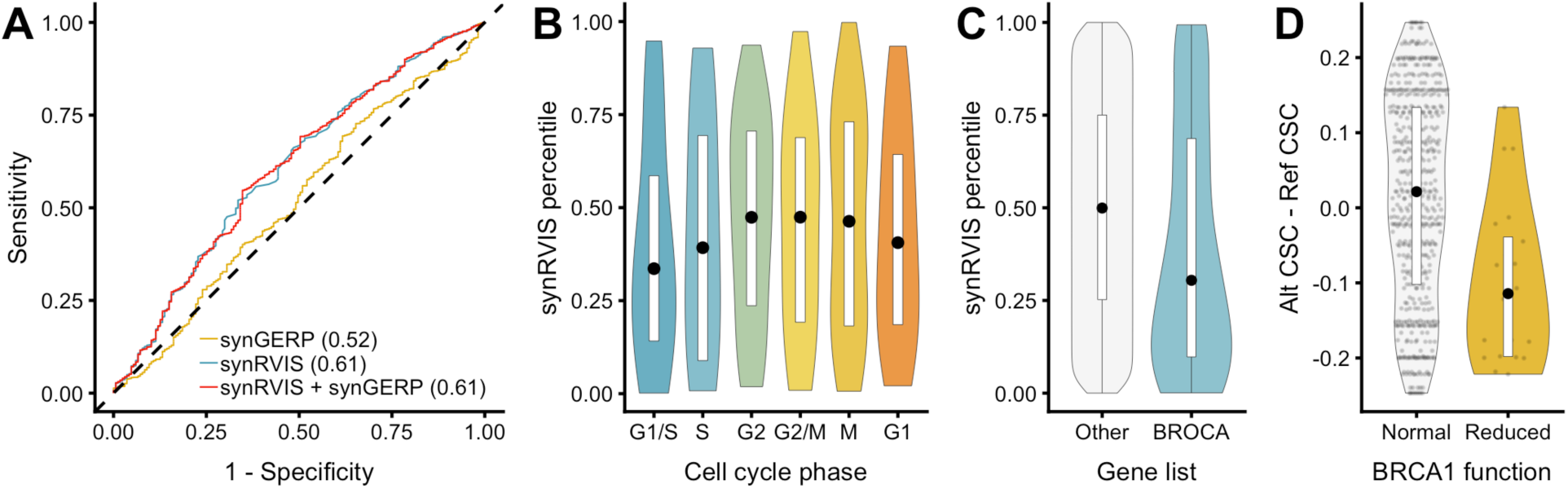
Intolerance of DNA damage response and cell cycle phase genes. **(A)** ROC curve illustrating the capacity of synGERP and synRVIS to predict DNA damage response genes. **(B)** synRVIS percentiles of genes periodically expressed in each cell cycle phase. **(C)** Distribution of synRVIS scores for genes contained in the BROCA Cancer Risk panel versus all other protein-coding genes. **(D)** Comparison of changes in CSC scores for synonymous variants in *BRCA1* that result in normal protein function versus those that reduce protein function.

tRNA levels also oscillate throughout the cell cycle, and genes that are expressed at different phases of the cell cycle have different codon usage (Frenkel-Morgenstern et al., 2012). In particular, tRNA expression levels are highest in the G2/M phase and lowest at the end of G1 phase. This coupling between tRNA expression and codon usage allows for cell cycle dependent oscillations in protein levels by ensuring that G2 phase genes are less efficiently translated during G1. Accordingly, we hypothesized that genes expressed during the G1 phase should be more intolerant to reductions in codon optimality than G2 genes. Strikingly, the synRVIS scores of these periodically expressed genes closely match the oscillatory changes in tRNA abundances; tolerance to reductions in codon optimality is lowest for G1/S expressed genes and increases stepwise by cell cycle stage, peaking for G2/M genes (**Fig 5B**). This finding not only supports previous observations about the codon usage patterns of cell cycle-related genes, but it provides the first direct evidence that these patterns are under selective constraint. synGERP scores did not display this pattern (**Fig S5A**), further suggesting that synRVIS may be more sensitive in detecting selection on genes that respond to tRNA availability.

The tRNA pool can also be dysregulated in disease, including certain cancers. Prior studies have found that oncogenic signaling upregulates particular tRNA species to enhance expression of pro-tumorigenic genes in breast cancer, ovarian cancer, and multiple myeloma (Gingold et al., 2014; Goodarzi et al., 2016; Pavon-Eternod et al., 2009; Winter et al., 2000; Zhou et al., 2009). We therefore hypothesized that cancer genes that are sensitive to shifts in tRNA abundances should be intolerant to changes in codon usage. To test this hypothesis, we compared the synRVIS and synGERP scores of hereditary breast and ovarian cancer genes included in the BROCA Cancer Risk Panel to all other protein-coding genes in the genome (Walsh et al., 2011; Walsh et al., 2010). This gene list includes 66 genes strongly implicated in hereditary breast and ovarian cancers. Accordingly, synRVIS but not synGERP scores were lower for these genes than the rest of the genes in the genome (synRVIS Wilcox *p* = .002, permuted *p* = .002; synGERP *p =* 0.80) (**Figs 5C** and **S5B**). Taken together, our results demonstrate the importance of codon usage in mediating gene expression under different physiological states.

### Synonymous variants that reduce codon optimality in *BRCA1* may abrogate protein levels

Collectively, our analysis suggests that synonymous mutations that alter codon optimality are under evolutionary constraint, meaning these mutations should have functional consequences. In particular, we expect that these variants may affect protein concentration by modulating mRNA translation and stability. To date, synonymous variants have been largely ignored in genetic disease association studies. However, synonymous mutations that reduce codon optimality in genes under strong selection could contribute to Mendelian disease. We have previously demonstrated that non-synonymous intolerance metrics, such as RVIS, play a pivotal role in discovering novel disease genes (Petrovski et al., 2013; Zhu et al., 2015). Our synRVIS and synGERP scores now provide, for the first time, a framework for identifying and prioritizing potential genes in which synonymous variants may also cause disease. Notably, genes with low synRVIS scores include genes such as *BRCA1* and *BRCA2*.

While functional impact of synonymous variants for most genes is unknown, we took advantage of a unique dataset in which CRISPR was used to perform saturation genome editing to assess the functional consequences of nearly all possible single nucleotide variants in the functionally critical RING and BRCT domains of *BRCA1* (Findlay et al., 2018). *BRCA1* ranked amongst the most highly intolerant genes (1^st^ percentile synRVIS, 13^th^ percentile synGERP) and loss of this protein predisposes women to breast and ovarian cancer (Hall et al., 1990; Kuchenbaecker et al., 2017). Thus, this dataset allows us to systematically answer the question of whether synonymous single nucleotide variants (SNVs) that reduce codon optimality significantly reduce BRCA1 dosage.

Findlay et al. introduced single nucleotide variants in the cell line *HAP1*, which is critically dependent on BRCA1 for cell survival. 11 days after introducing the mutations, they sequenced the line to gauge the frequency of each variant in the cell population, as variants that reduce BRCA1 expression or function result in cell death. These frequencies were converted to a continuous score and variants that were less frequent in the population than average were presumed to have resulted in reduced BRCA1 activity. They also performed measured expression of *BRCA1* to assign RNA scores to reflect each variant’s effect on gene expression.

Of roughly 500 introduced synonymous mutations in *BRCA1*, 19 were associated with scores that signified reduced BRCA1 activity. We hypothesized that reduced function scores for synonymous variants were likely indicative of decreased expression and/or translation. For each synonymous variant, we calculated the difference between the CSC value of the alternate and reference alleles, such that negative changes signify reductions in codon optimality. Accordingly, the 19 synonymous mutations associated with reduced BRCA1 activity were significantly more likely to attenuate codon optimality (Wilcox *p* = .001) (**Fig 5D**). We also calculated the correlation between the RNA and function scores and the difference in CSC values for all synonymous variants assayed (**Fig S5C, D**). We found that changes in codon optimality significantly correlated with *BRCA1* function scores (Pearson’s *r* = .27, *p* = 3 × 10^−11^) and RNA scores (Pearson’s *r* = 0.15, *p* = 4.8 10^− 4^). Although these correlations are modest, they suggest that at least a fraction of variants that reduce codon optimality may have functional consequences in BRCA1, presumably via modulation of translation and/or mRNA stability. We note that there are other potential mechanisms by which these variants could functionally impact *BRCA1*, including via modulation of splicing enhancers. Therefore, further molecular studies are required to elucidate the precise functional consequences of attenuated codon optimality in *BRCA1*.

Nonetheless, these results imply that some synonymous variants that affect codon usage can result in effect sizes that could cause Mendelian disease. synRVIS thus provides an initial framework for identifying putatively pathogenic synonymous mutations in the interpretation of human genomes, as mutations in genes most intolerant to synonymous variation are more likely to be pathogenic. Importantly, 3 of the 19 synonymous variants that reduce BRCA1 function appear in gnomAD, indicating that individuals do in fact harbor potential disease-causing synonymous variants that may be overlooked in standard carrier screens.

## Discussion

Through comprehensive analyses, we demonstrate the role of natural selection in optimizing the codon content of the human genome. First, we show that synonymous mutations that reduce codon optimality appear at lower allele frequencies in the human population than neutral variants and variants that increase codon optimality. Supporting this result, we find that synonymous sites in optimal codons tend to be more strongly phylogenetically conserved across the mammalian lineage. We introduce two per-gene intolerance scores, synRVIS and synGERP, which assess the strength of selective constraint on synonymous variation in each protein coding gene. synRVIS detects human-specific selection against variants that reduce codon optimality, whereas synGERP reflects the phylogenetic constraint of fourfold degenerate sites in a given gene. We find that these scores predict dosage-sensitive genes, emphasizing the importance of codon usage in mediating protein concentration.

Recent studies have revealed that synonymous codon usage serves as a secondary genetic code that guides translation efficiency and mRNA stability in human cells (Forrest et al., 2018; Wu et al., 2019). In particular, translation elongation rate, which is partially a function of tRNA abundance, is posited to impact mRNA degradation rate. Despite these molecular consequences, population geneticists have argued that the effect size of any single synonymous SNV would be too small to be selected against in the human population (Chamary et al., 2006). Our results cast doubt on this assumption in two ways. First, the allele frequency distributions illustrate that there are genome-wide signatures of selection against reductions in codon optimality. This finding shows that some synonymous mutations are of large enough effect to be selected against even in the context of the small human effective population size. Second, we demonstrate that some codon optimality-reducing SNVs in *BRCA1* can significantly attenuate protein activity, potentially via reduced mRNA stability and translation. These findings are consistent with a handful of other studies that have implicated synonymous SNVs in human disease (Dershem et al., 2018; Hunt et al., 2014; Kimchi-Sarfaty et al., 2007). In sum, the effect sizes of synonymous mutations have been historically underestimated. A fraction of these variants can significantly reduce protein concentration to the same extent as loss-of-function variants (Dershem et al., 2018) and likely represent an important, albeit typically ignored, source of Mendelian disease. Furthermore, even synonymous SNVs that only modestly reduce protein output could play a significant role in both modifying mendelian disease and in the genetic architectures of complex traits, which seem to be driven by the cumulative effect of many small effect size variants (Boyle et al., 2017; Yang et al., 2010).

Our results support the functional relevance of translational regulation of gene expression. Consistent with the effects of translational efficiency on protein output and mRNA stability, we find that dosage-sensitive and loss-of-function depleted genes tend to be more intolerant to synonymous variation. However, one limitation of our study is that the calculation of synRVIS relies on codon usage metrics derived from a single cell type whereas tRNA expression varies widely by tissue (Dittmar et al., 2006). synGERP, on the other hand, does not rely on codon usage scores, but is less sensitive to detecting constraint on potential human-specific tRNA expression dynamics.

We found that some loss-of-function depleted genes involved in neurodevelopment were in fact very tolerant to reductions in codon optimality. Intriguingly, a recent study found that codon optimality is attenuated in genes expressed in the developing *Drosophila* nervous system (Burow et al., 2018). This reduced optimality mitigates the effect of codon content on mRNA stability, thereby allowing *trans*-acting factors, such as RNA-binding proteins and microRNAs, to exert greater influence over mRNA decay in the developing nervous system. If this phenomenon exists in humans, it could explain our observation that some LOF-depleted genes are tolerant to changes from optimal-to-nonoptimal codons. Additionally, because tRNA expression is likely markedly different in the brain (Bazzini et al., 2016; Bornelov et al., 2019; Dittmar et al., 2006), synRVIS may be limited in detecting intolerance of neurodevelopmental genes due to its reliance on HEK293T-derived codon stability coefficients. Both of these hypotheses could explain synGERP’s improved ability to predict dosage sensitive genes since synGERP detect constraint on binding sites for *trans-*acting factors and does not rely on CSC in its calculation. The role of codon usage in the unique translational demand of neurons remain an outstanding question. Furthermore, understanding the relationship between tissue-specific codon usage, intolerance, and mRNA decay programs stands as an important goalpost for future studies.

Strikingly, we not only found a correlation between the strength of selection on codon optimality and disease relevant genes, but we also found a relationship with the tRNA abundance patterns that prevail when specific genes are expressed. Specifically, changes in tRNA abundance can modulate protein expression in response to different cellular states, including cell cycle stage, disease, and stress. Previous studies have demonstrated that cellular tRNA levels are reduced in response to DNA damage and during the G1 phase of the cell cycle (Frenkel-Morgenstern et al., 2012). Accordingly, we illustrate that intolerant synRVIS genes are enriched for genes involved in these cellular pathways. synGERP is unable to predict these genes, perhaps implicating a role of human-specific selection on codon optimality in these pathways. We note that tRNA dysregulation also underpins the pathogenesis of other non-cancerous conditions, including some immunodeficiency and neurological disorders (Morita et al., 2013; Piccirillo et al., 2014; Tahmasebi et al., 2018). Therefore, future work focused on determining potential interspecies variation in dynamic tRNA expression will be crucial in determining whether non-human disease models accurately represent diseases characterized by translational deregulation.

Collectively, our results suggest that codon usage can significantly impact biological traits and may play an underappreciated role in human disease. Just as previously developed intolerance scores have transformed our ability to identify diseause-causing non-synonymous variants (Petrovski et al., 2013; Zhu et al., 2015), synRVIS and synGERP will aid in identifying synonymous variants that drive human traits. We note that synRVIS critically depends on codon usage metrics and the number of individuals sequenced in the reference cohort. Therefore, our resolution to detect intolerance to synonymous variation in the human genome will improve with tissue-specific codon stability coefficients and increased numbers of sequenced individuals.

## Supporting information

All supplemental figures

## Acknowledgements

We wish to thank many people for very helpful discussions or comments, including Slavé Petrovski, Chirag Vasavda, and Gundula Povysil. We also thank Chirag Vasavda, Brian Khoe, Daniel Zhang, and Xinchen Wang for feedback on figure design.

## Author contributions

R.S.D., A.M.M., and D.B.G. conceived of the study and wrote the manuscript. R.S.D. and B.R.C. performed analyses.

## Methods

### Sequence data

We used summary level allele frequency data from the BRAVO TOPMed database (TOPMed Freeze 5, https://bravo.sph.umich.edu/freeze5/hg38/) and from the gnomAD exome database (release 2.0.2., https://gnomad.broadinstitute.org/downloads). The TOPMed database contains roughly 463 million variants derived from 62,784 whole genomes and the gnomAD exome database contains roughly 15 million variants from 123,136 whole exomes.

We mapped TOPMed variants from hg38 to hg19 using the LiftoverVCF tool in Picard tools (http://broadinstitute.github.io/picard/, 2019) (v2.9.0). We then annotated both the TOPMed and gnomAD VCFs using Variant Effect Predictor (VEP), version 84 (McLaren et al., 2016). We used the VEP “--pick_allele” option with the following order: “rank, canonical, appris, tsl, biotype, ccds, length” to annotate each variant with its most damaging possible effect across all transcripts.

We then filtered each VCF file to contain only variants annotated as “PASS” and removed all variants occurring in repeat regions, as identified by RepeatMasker, version 4.0.5 (Smit, 2013). To exclude variants that are expected to disrupt canonical splice sites, we removed all variants occurring within 10 intronic nucleotides and 3 exonic nucleotides of exon-intron boundaries in all Ensembl v75 transcripts. We additionally filtered the gnomAD VCF to only retain variants with at least 10-fold coverage in at least 85% of individuals.

### Codon usage metrics

We used two scores for assessing codon usage: the codon stability coefficient (CSC) and the relative synonymous codon usage (RSCU). We obtained CSC scores derived from HEK293T cells (Wu et al., 2019). Wu et al. also calculated CSC scores for other cell lines, including HeLa and RPE cells, but these scores were very strongly correlated with the HEK293T scores. The CSC represents the Pearson correlation between the frequency of the codon in each transcript and the associated half-life. We classified codons with a CSC values greater than 0 as “optimal” and codons with CSC values less than 0 as “non-optimal.”

As an orthogonal measure of codon usage, we calculated RSCU scores for each codon (Sharp and Li, 1986). For each codon in each canonical transcript (as defined by Ensembl v75), we calculate the ratio of the observed number of codons to the expected number for a given amino acid. Specifically, for an amino acid *i*, the RSCU score of its *j^th^* amino acid is defined as:

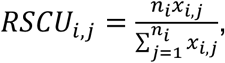

where *n_i_*denotes the number of synonymous codons for that amino acid. When using RSCU to assess codon optimality, we annotate codons with a value less than 1 as “non-optimal” and greater than 1 as “optimal.” We chose to calculate gene-specific rather than genome-wide RSCU scores, reasoning that gene-specific scores should more adequately reflect tissue-specific sources of constraint.

### Site frequency spectrum analyses

We performed all SFS analyses using the filtered TOPMed allele frequency data. We adapted an approach previously employed in *Drosophila* studies to compare selection on synonymous variation with putatively neutral variants (Lawrie et al., 2013; Machado et al., 2017). Specifically, we matched each observed synonymous variant occurring at fourfold degenerate sites with intronic variants occurring within 10,000 basepairs. We required matched variants to have the same ancestral allele, and in an additional analysis, we required matched variants to also have the same neighboring 5’ and 3’ nucleotides. We matched blind to the direction or strand, such that synonymous mutations were allowed to pair with forward, reverse, and reverse complement intronic sequences. For all SFS analyses, we only considered synonymous variants occurring at fourfold degenerate sites.

We folded all allele frequencies: if the alternate allele frequency was greater than 50%, we subtracted it from 100%, meaning the minor allele frequency is always less than or equal to 50%. We then used a two-tailed t-test to determine whether SFS distributions were significantly different (Harpak et al., 2016; Keinan et al., 2007). As noted by Keinan et al, this test is conservative since it reflects significant deviation in the mean minor allele frequencies rather than other differences in the shape of the distribution (2007).

### Comparing phylogenetic conservation at synonymous and intronic sites

We used a custom script to annotate the TOPMed variants with GERP++ scores (Davydov et al., 2010). To assess phylogenetic conservation on codon usage, we compared the GERP++ scores of the reference alleles of the synonymous and intronic variants included in the SFS analyses. Because only a fraction of fourfold degenerate sites actually harbors a variant in TOPMed, we also compared the correlation between CSC and GERP++ at all fourfold degenerate sites in the genome. To mitigate confounding due to conservation at splice sites, we excluded codons occurring at exon-intron boundaries in all Ensembl v75 transcripts.

### Deriving synRVIS

Using aggregated allele frequency from gnomAD exomes (Karczewski et al., 2019), we defined Y as the total number of common (MAF > 0.5%) synonymous O → NO SNVs in a gene and X as the total number of synonymous SNVs occurring in a gene. We then regressed Y on X and defined synonymous RVIS (synRVIS) as the studentized residual for each gene. The resulting regression line accounts for genic mutation rates, sequence context, and gene size while predicting the expected number of common synonymous variants that result in a non-optimal change. We explored the behavior of the score when we used alternative MAF cutoffs of 1% and 0.1% for defining common variants on the Y-axis and found that these scores strongly correlated (Pearson’s *r* = 0.89 and *r* = 0.74, respectively) (**Fig S3A-B**). We also found strong correlation when we used RSCU instead of CSC to define codon optimality (*r* = 0.63) (**Fig S3C)**.

### Calculating synGERP

We defined the synGERP score as the average GERP++ score (Davydov et al., 2010) of all fourfold degenerate sites in a given gene. We excluded all codons immediately adjacent to exon-intron junctions in all Ensembl v75 transcripts to mitigate confounding due to conservation at canonical splice sites.

### Gene set enrichment tests

We used logistic regression models to determine the ability of synRVIS and synGERP to predict 360 dosage sensitive genes contained in the ClinGen Genome Dosage Map (http://www.ncbi.nlm.nih.gov/projects/dbvar/clingen/) and 178 DNA damage response genes (http://sciencepark.mdanderson.org/labs/wood/dna_repair_genes.html). We calculated receiver operating characteristic (ROC) curves using the pROC package in R (Robin et al., 2011). For the BROCA cancer risk panel genes (https://testguide.labmed.uw.edu/public/view/BROCA), we opted to perform a Mann Whitney U test to compare the intolerance of these genes versus all other genes in the genome rather than evaluate the ROC, given the small sample size of the gene list (n=66). We additionally performed a permuted Mann Whitney U test for this particular enrichment test. Specifically, we first computed the actual Mann Whitney U p-value of the observed data. We then randomly permuted the labels of the data and computed additional p-values 1,000 times. We defined the permuted p-value as the proportion of permuted p-values less than or equal to the actual p-value derived from the original, unpermuted dataset.

We also compared the distribution of synRVIS and synGERP to LOEUF (https://gnomad.broadinstitute.org/downloads), a metric that assesses the observed over expected ratio for loss-of-function variants in the gnomAD database. Specifically, we computed the median LOEUF score per synRVIS and synGERP decile. We also assessed the median synRVIS and synGERP percentile of genes dynamically expressed during the cell cycle, as identified by CycleBase (https://cyclebase.org/CyclebaseSearch). All gene lists are available in **Table S2**.

### GO Enrichment

We performed GO enrichment tests of genes falling below the 25^th^ percentile in synRVIS or synGERP to identify classes of genes most intolerant to synonymous variation. We also performed enrichment tests of synRVIS tolerant but LOEUF intolerant genes. We defined these genes as genes above the 75^th^ percentile in synRVIS, but below the 25^th^ percentile of LOEUF scores based on our observation in **Figure 4B**. To perform the enrichment test, we used the PANTHER GO Enrichment Analysis Tool (http://geneontology.org), using the PANTHER GO-Slim Biological Process annotation set. P-values were computed using Fisher’s Exact Test and corrected via False Discovery Rate. We defined corrected p-values < .05 as significant. The full lists of significant GO enrichment results are available in **Table S3.**

### BRCA1 function score evaluation

We used VEP to annotate the resulting codon changes from synonymous variants assayed by Findlay et al (2018). We then annotated the reference and alternate codons of each variant with their CSC values and removed all variants identified as splice region variants by VEP or occurring within 3 basepairs of exon-intron junctions (**Table S4**). We annotated variants with function scores less than −0.748 as variants that reduced BRCA1 function. To quantify codon usage changes, we defined ΔCSC as the difference between the alternate CSC value and the reference CSC value for each variant.

### Data visualizations

All plots were generated in R using ggplot2 (Wickham, 2016). Figure 1A was created with BioRender (https://biorender.com). Color palettes for plots were derived from the wesanderson R package (https://github.com/karthik/wesanderson).

## Supplementary tables

Table S1: List of genes with their synRVIS and synGERP scores

Table S2: Gene lists used for enrichment tests

Table S3: GO enrichment results

Table S4: Annotated *BRCA1* variants from Findlay et al. (2018).

