## Supplementary material for "Natural selection shapes codon usage in the human genome": All supplemental figures

### Supplementary figures

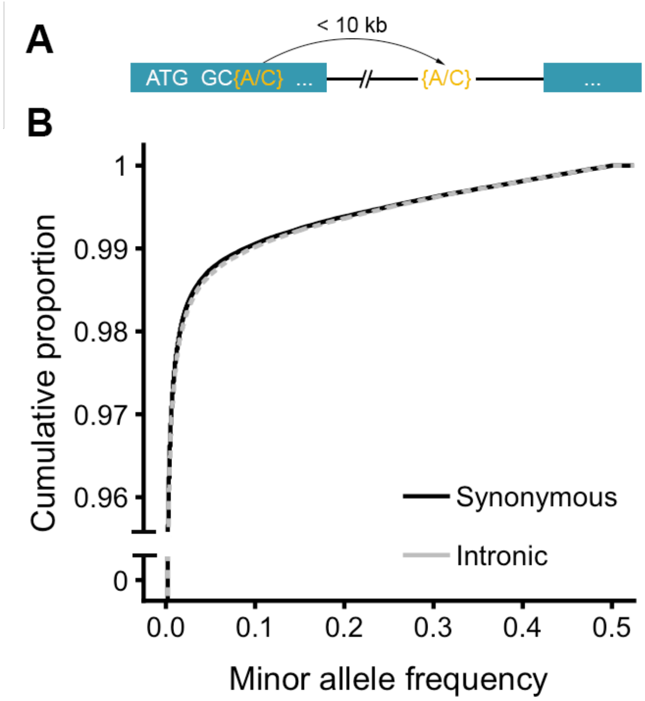


**Figure S1. Demonstration of variant matching scheme and baseline SFS. (A)** Illustration of variant matching scheme for TopMed variants. Each observed synonymous variant was matched to an observed intronic variant within 10kb and with the same reference and alternate allele. We excluded all variants occurring in the first and last codon of an exon and intronic variants within 10 basepairs of splice junctions. **(B)** Site frequency spectrum of synonymous and intronic variants without accounting for codon bias.

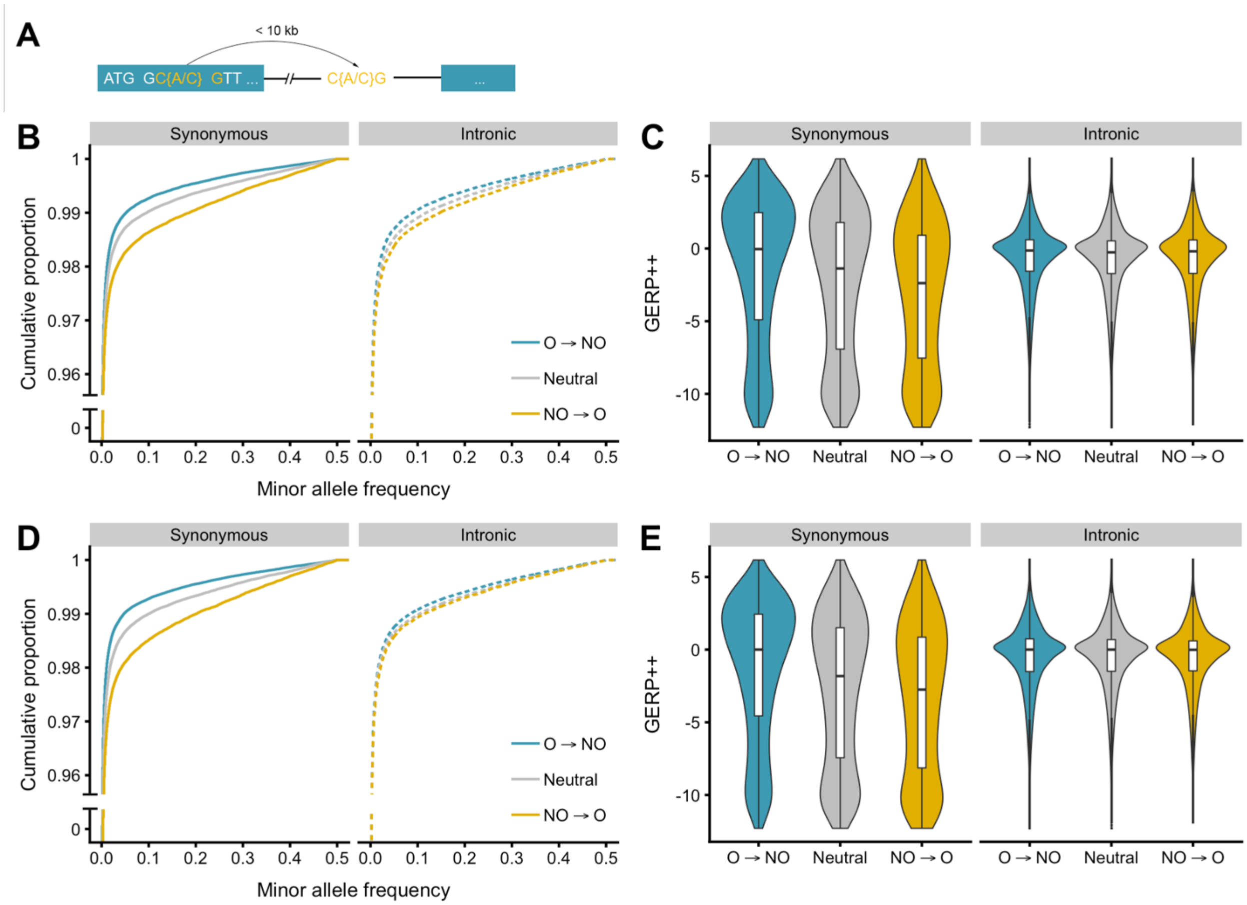


**Figure S2. SFS of synonymous and intronic variants matched for 5' and 3' nucleotide content (A)** Illustration of variant matching scheme for TopMed variants that matches synonymous and intronic variants that share the same major and minor allele as well as the same neighboring 5' and 3' nucleotide. **(B)** Site frequency spectrum of matched variants. T-test p-values: Syn O → NO vs syn neutral ($p=3.2x10^{-34}$); syn O → NO vs intronic O->NO ($p=5.4x10^{-27}$); syn NO → O vs intronic NO → O ($p=6.2x10^{-4}$); and syn NO → O vs syn neutral ($p=9.1x10^{-16}$) **(C)** GERP++ distributions of the reference alleles of the matched variants. **(D)** SFS of the original matched synonymous and intronic variants using RSCU-defined codon optimality. **(E)** GERP++ distribution of reference alleles of the RSCU-annotated variants.

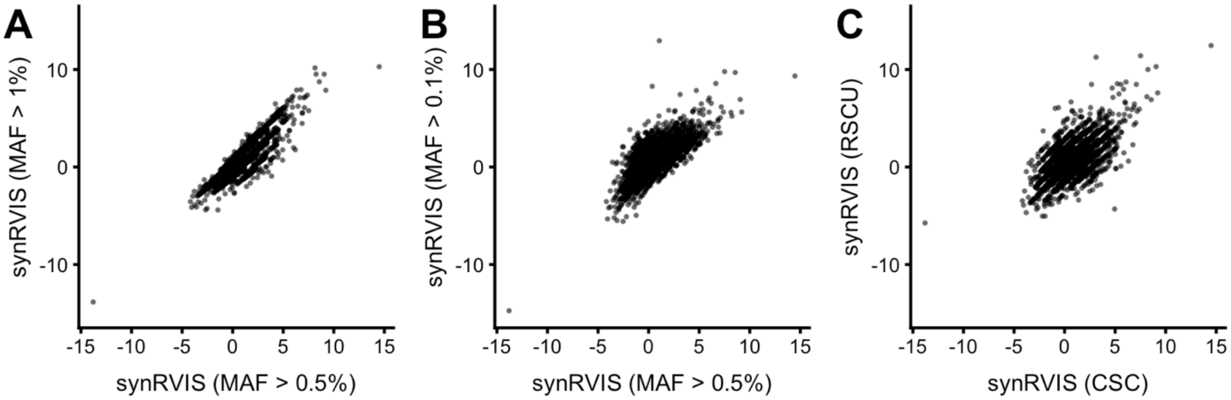


**Figure S3. Scatter plots of alternative synRVIS derivations. (A)** Comparison of synRVIS scores calculated using a 1% MAF cutoff rather than 0.5% MAF cutoff for defining common optimal to non-optimal codons (Y). **(B)** Comparison of using a MAF cutoff of 0.1% rather than 0.5% for (Y). **(C)** Comparison of CSC-defined codon optimality versues RSCU-defined codon optimality (MAF cutoff of 0.5% for both).

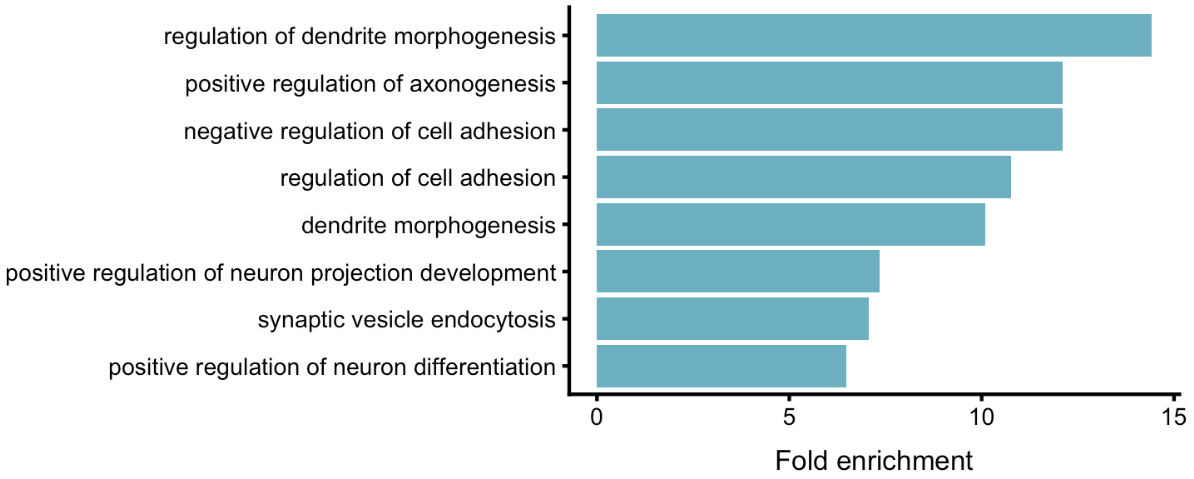


**Figure S4: GO enrichments of genes tolerant to synonymous variation but intolerant to loss-of-function variation.** Top GO categories enriched for genes below 25$^{th}$ percentile LOEUF and top 25$^{th}$ percentile synRVIS.

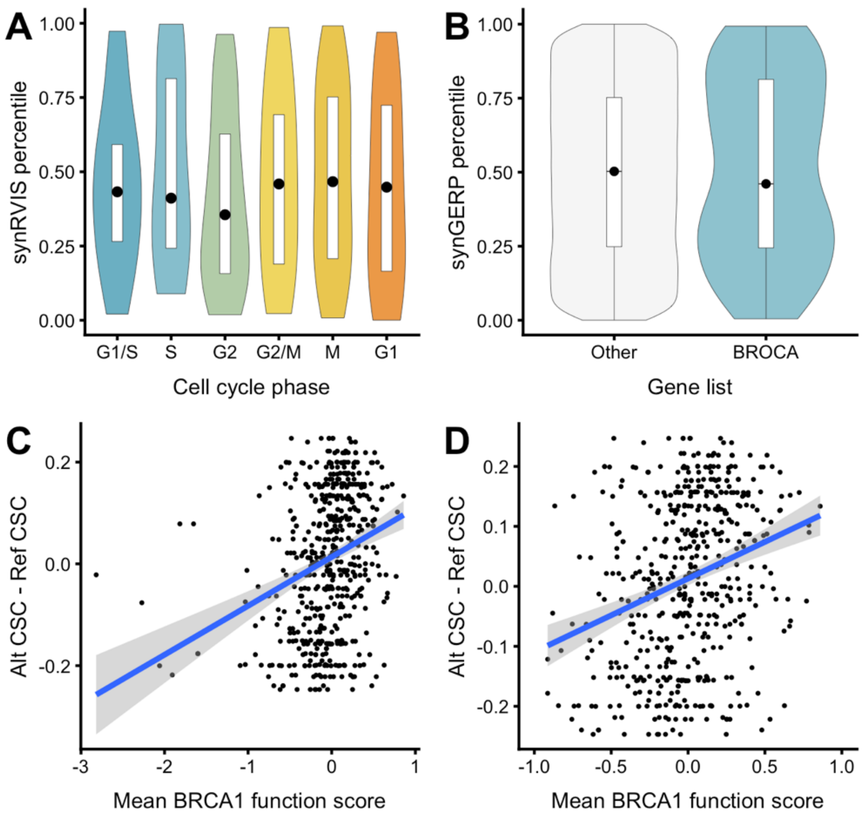


**Figure S5. synGERP distributions of cell cycle expressed genes and BROCA list genes. (A)** synGERP distributions of genes periodically expressed in the cell cycle. **(B)** synGERP distribution of genes contained in the BROCA panel versus all other protein coding genes. **(C)** Scatter plot of CSC scores and function scores for synonymous variants in *BRCA1*. **(D)** Same as (C) with outliers removed.
